# Interactome Analysis of the CC2D1A Scaffold Reveals Novel Neuronal Interactions and a Postsynaptic Role

**DOI:** 10.1101/2025.06.26.661826

**Authors:** Abigail T. Heller, Aniket Bhattacharya, Haorong Li, Luka Turkalj, Shruthi Thiyagarajan, Emma Suzuki, Adele Mossa, Haiyan Zheng, Ling Hao, M. Chiara Manzini

## Abstract

Loss of the protein scaffold Coiled-coil and C2 domain containing 1A (CC2D1A) leads to intellectual disability (ID), autism spectrum disorder (ASD), and other neurodevelopmental presentations in humans. CC2D1A interactions have been studied in different cell lines proposing diverse roles in endolysosomal maturation and intracellular signaling, but the composition and functional mechanisms of the CC2D1A interactome remain poorly understood, especially in the brain. We performed comprehensive proteomic analyses to characterize CC2D1A binding partners, first comparing immunoprecipitations with three different anti-CC2D1A antibodies in HEK293 cells and then probing the mouse hippocampus.

In HEK cells, Gene Ontology (GO) analysis revealed broad interaction networks in the nucleus, mitochondrion, and cytoplasmic vesicles sharing functions in organelle organization, vesicle mediated transport, and protein metabolism. These are unified by the best characterized CC2D1A interactor, the ESCRT III component CHMP4B, and define a pleiotropic role for CC2D1A in membrane trafficking and protein homeostasis. In the hippocampus, using stringent criteria and additional controls, including a *Cc2d1a* hypomorph mouse line, we identified 10 high-confidence interactors in addition to CHMP4B (TNIK, G3BP2, CEP135, MAPKAP1, SHFL, PPT1, PNKD, VAMP5, and PPP6R2) revealing roles for RNA regulation and synaptic function. The HEK studies had also pointed to CC2D1B, the only paralog of CC2D1A, as an interactor. We confirmed that not only the two proteins can bind in the brain, but also localize in different synaptic compartments, showing that CC2D1A is uniquely enriched in the post-synapse. This supports a unique function of CC2D1A in regulation of synaptic transmission that could explain the more severe cognitive deficits in humans and mice upon its loss. To our knowledge these findings provide the most comprehensive characterization of the CC2D1A interactome to date, elucidating novel, multifaceted, and dynamic cellular functions, providing potential implications for its role in neurodevelop-mental disorders.

## INTRODUCTION

Coiled-coil and C2 domain containing 1A (CC2D1A; MIM*610055, UniProtKB Q24K25) is a multifunctional signaling scaffold that is mutated in rare, autosomal recessive form of intellectual disability (ID), often comorbid with autism spectrum disorder (ASD) (1-8). The 951 amino acid protein has a unique structure with four conserved DM14 (*Drosophila melanogaster* 14) domains, two coiled-coil domains, and one C2 domain. DM14 domains are only present in metazoans and are exclusive to the CC2D1 protein family. These proteins are present as a single gene in invertebrates with *Drosophila* Lgd (Lethal (2) giant disks) as the best studied ortholog, while two paralogs with similar organization - CC2D1A and CC2D1B – are found in vertebrates (9). The DM14 repeats are short helical hairpins that facilitate protein-protein interactions (10). The C2 domain is a Ca^2+^-dependent membrane-targeting domain shared across multiple proteins (11), and in case of CC2D1A, it has been proposed to tether the CC2D1A complex to the phospholipid membrane to mediate signaling activation (12, 13).

Known interaction partners of CC2D1A have mostly been identified through one or two-hybrid assays and converge on endosomal trafficking and intracellular signaling. Various studies pointed to the CC2D1 proteins as critical regulators of the ESCRT (endosomal sorting complex required for transport) pathway by facilitating CHMP4B polymer assembly to regulate the dynamics of membrane scission in endolysosomal sorting of transmembrane receptors (9, 10, 14-17) as well as HIV budding (14). CC2D1A was also found to act as a scaffold for signaling ranging from facilitating PDK1/AKT complex formation to regulate AKT kinase activity (18), endosomal targeting of regulators of antiviral immunity such as RIG-I, TLR3, and TLR4 (16, 19), and modulating ERK signaling via EGFR (16). Numerous other binding partners such as steroid receptor co-activator 2 (SRC2), Tank-binding kinase 1 (TBK1), EGFR have also been identified (18, 20). In addition, CC2D1A binds to phosphodiesterase 4D (PDE4D), a cAMP scavenger, repressing its activity to control cyclic AMP (cAMP) levels and regulate cAMP-dependent protein kinase (PKA) activity (21-23). In both *CC2D1A* knock-out cells and animal models, PDE4D is constitutively phosphorylated and hyperactive, leading to a reduction in cAMP-PKA signaling (12, 22, 23). Finally, CC2D1A has also been reported to function as a transcription factor in serotonergic raphe neurons where it represses 5-HT1A receptor transcription (24). While these findings indicate that CC2D1A is a multifunctional protein with roles in intracellular signaling (2, 16, 18, 19, 23) and endosomal trafficking (9), the underlying molecular mechanisms of neurodevelopmental deficits caused by *CC2D1A* loss of function remain poorly understood.

ASD and ID risk genes show remarkable genetic heterogeneity and can act on a variety of molecular pathways regulating synaptic transmission and intracellular signal transduction (25-27). CC2D1A has been involved in multiple signaling cascades that are known to regulate neuronal development, differentiation, synaptic maturation and plasticity and mirrors this pleiotropy (2, 12, 13, 18, 22, 23, 28, 29). Previous proteomic work attempting to define the CC2D1A interactome used an antibody with unclear specificity (12). To address this limitation, we designed a highly controlled experiment comparing three different commercial antibodies raised against CC2D1A and compared interactors in non-neuronal cell lines (HEK293) and in the brain. While most biallelic loss of function variants in *CC2D1A* lead to ID and ASD supporting a critical role in brain development (1-8, 30-32), CC2D1A is ubiquitously expressed and some variants have been recently shown to cause broader impacts with ciliopathy and heterotaxy (6, 33). In addition, multiple studies revealed non-neuronal function that may be redundant with CC2D1B (9, 14, 16, 34). Given such functional diversity, we hypothesized that the CC2D1A interactome will vary across tissue types. Here, we report the interactome in both a human cell line (HEK293) as well as mouse hippocampus. We found that CC2D1A can be deployed to different subcellular locations in conjunction with CHMP4B and other ESCRT components to interact with proteins involved in organelle organization, protein trafficking, and proteostasis. In the brain, we identified novel interactions with signaling proteins and CC2D1A enrichment over CC2D1B at the post-synaptic density which may explain its unique involvement in cognitive dysfunction.

## METHODS

### Animals

All mouse experiments were carried out as approved by the Rutgers University Institutional Animal Care and Use Committee (IACUC) and following National Institutes of Health guidelines. All animals used were on a C57BL/6J background (Charles River Laboratories). The *Cc2d1a-V5HA* (hereafter 1aKD or VH/VH) mouse line was generated by Clustered Regularly Interspaced Short Palindromic Repeats (CRISPR) insertion of a 78 base pair (bp) tag containing both V5 and HA sequences in the 3’ end of the *Cc2d1a* gene (**Supplementary Fig.S1**). The insertion and orientation of this tag was verified by sequencing pups that inherited the tag through the germline and by Western blotting for an anti-HA antibody (Invitrogen:26183). Mice were genotyped using PCR with primers (forward 5’-CCT GGT GGA GAG CGA GGT AA-3’, reverse 5’-CGC TGA TTG TCT GCT-3’ bp).

### Protein Lysate Preparation

Human Embryonic Kidney (HEK293) cells were purchased from American Type Culture Collection (CRL-1573, ATCC Manassas, VA) and sub-cultured as per standard protocols (3). Approximately 90% confluent cultures were washed once with 2 mL of cold 1X phosphate buffered saline (PBS), following media removal, and lysed in 1 mL of freshly prepared 0.1% sodium dodecyl sulphate (SDS) in 1X PBS. Both PBS and the lysis buffer were supplemented with 1:500 each of protease (Sigma-Aldrich:P8340) and phosphatase inhibitors (Sigma-Aldrich:P0044). Lysates were then clarified by passing through a 28G1/2 1cc syringe and centrifuged at 15,000 rpm for 20 minutes at 4°C. The supernatant was collected in a fresh tube, quantified using BCA assay (G-Biosciences:786-571) and stored frozen in -80°C until immunoprecipitation.

Hippocampi were dissected from two-month-old mice in ice-cold 1X PBS, flash-frozen in liquid nitrogen, and stored at -80°C until lysate preparation. Lysates were prepared in non-denaturing lysis buffer (100 mM NaCl, 20 mM Tris-HCl pH 7.4, 1 mM ethylenediaminetetraacetic acid (EDTA) solution supplemented with 1:500 diluted protease and phosphatase inhibitors) by gently homogenizing the frozen tissue with disposable plastic pestles (VWR:66001-104). Following a 10-minute incubation on ice, the lysates were clarified by centrifugation at 15,000 rpm for 20 minutes at 4°C, the supernatant was collected and quantified using BCA assay (G-Biosciences:786-571). Lysates from 2-3 mice (4-6 hippocampi) were pooled to obtain sufficient protein for each experiment. For each set, 2-3 mg protein were used in total, which was then evenly divided between either the anti-CC2D1A antibody or IgG-coated beads immunoprecipitation tubes after preclearing.

### Immunoprecipitation (IP)

Protein lysates were pre-cleared with control resin beads (Pierce:26150) that had been prewashed thrice with 1 mL cold lysis buffer and then incubated with 2 mg/mL of lysate for at least 1 hour at 4°C, under gentle rocking. Antibodies were crosslinked on A/G protein agarose beads (Pierce:20423). Beads were prewashed thrice with 1 mL phosphate buffer (0.1 M sodium phosphate dibasic, 0.1 M sodium phosphate monobasic, pH 7.2) and incubated with CC2D1A or IgG antibodies for 30 minutes at room temperature with end-over rolling using 15 µg of each antibody for mass spectrometry and 10 ug for western blot validation. Anti-CC2D1A antibodies used were a rabbit monoclonal (Abcam:ab191472), a mouse polyclonal (Abcam:ab68302), or a rabbit polyclonal (Bethyl:A300-285A). Matching negative controls were either an anti-rabbit IgG (EMD Millipore:12-370) or an anti-mouse IgG (EMD Millipore:12-371) depending on the host species of the IP antibody. Excess primary antibodies were washed off thrice with 1 mL phosphate buffer, and antibody-coated beads were incubated with 50 µl of 450 µM DSS crosslinker (ThermoScientific:1863418) for 30 minutes of room temperature with end-over rolling. Cross-linked beads were then washed thrice with 1mL phosphate buffer. For IP, 1 mg of precleared lysate was used per condition with either anti-CC2D1A antibody or IgG-coated beads for overnight incubation at 4°C with gentle rocking. The next day, the beads were spun at 10,000 rpm and the flowthrough was collected. The beads were washed five times with cold lysis buffer. The complexes were eluted from the beads in 0.1 M glycine (pH 2.5), with agitation (10,000 rpm) at 56°C for 10 minutes, then neutralized with Tris-HCl pH 8, and prepared for western blot. For MS-MS, the beads were additionally subjected to two extra washes in a wash buffer (100 mM NaCl, 20 mM Tris-HCl pH 7.4) to remove EDTA, pelleted, frozen, and submitted to the Rutgers Proteomics Core. IPs were performed in duplicates on HEK293 cells for each anti-CC2D1A antibody. Mouse hippocampal IPs were performed in duplicate for age-matched WT males and females and then analyzed together with one male and two female samples for both WT and 1aKD mice. Hence, a total of 3 WT males and 4 WT females were compared across two independent LC-MS/MS runs. Protein digestion was conducted on beads. 0.2 μg of trypsin in 20 ml 50 mM NH_4_HCO_3_ was added to the IP beads and incubated at 37°C for 4 hours. Another 0.2 μg of trypsin was added and incubated overnight at 37°C. The solution was separated from the beads, and pH was adjusted to 3 with 10% formic acid. The sample was desalted with Stage-tip before LC-MS/MS (35).

### Liquid chromatography-tandem mass spectrometry (LC-MS/MS)

Samples were analyzed on a Dionex Ultimate 3000 RLSCnano System interfaced with an Orbitrap Eclipse Tribrid mass spectrometer (ThermoFisher). Samples were loaded on to a fused silica trap column (Acclaim PepMap 100, 75 μm x 2 cm, ThermoFisher). After washing for 5 minutes at 5 µl/min with 0.1% TFA, the trap column was brought in-line with an analytical column (Nanoease MZ peptide BEH C18, 130A, 1.7 μm, 75μm x 250 mm, Waters) for LC-MS/MS. Peptides were separated at 300 nL/min using a segmented linear gradient 4-15% B in 30 minutes (where A: 0.2% formic acid, and B: 0.16% formic acid, 80% acetonitrile), 15-25% B in 40 minutes, 25-50% B in 44 minutes, 50-90% B in 11 minutes, and 4% for 5 minutes. MS1 scanned from *m/z* 350-1600 with a resolution of 120,000, an automatic gain control (AGC) target of 1E6, a maximum injection time of 100 ms. Top 3 seconds data-dependent acquisition was conducted with a dynamic exclusion of 60 seconds, a quadrupole isolation window of *m/z* 1.4, MS/MS resolution of 30,000, an AGC target of 1E5, and a normalized collision energy of 30%. The MS/MS scan range were determined by charge state of the parent ion, but lower limit was set at 100 amu.

### Proteomics data analysis

LC-MS/MS raw files were analyzed using the Thermo Fisher Proteome Discoverer software (2.4.1.15) with the SequestHT search engine. Reviewed Swiss-Prot *Homo sapiens* (20,328 protein entries, downloaded December 5, 2022) or *Mus musculus* (17,114 protein entries, downloaded December 5, 2022) database was used for human and mouse protein identifications, respectively. Mouse tissue or cell culture specific contaminant FASTA libraries were used to mark and remove contaminant proteins (available to download at https://github.com/HaoGroup-ProtContLib, (36)). Precursor mass tolerance was set to 25 ppm. Fragment mass tolerance was set to 0.02 Da. False discovery rate cutoff was set at 1% for protein and peptide spectral match identifications. Trypsin was selected as the enzyme with four maximum mis-cleavages allowed. Cysteine alkylation was included as a fixed modification. Methionine oxidation and acetylation of protein N-terminus were included as variable modifications. Mass tolerance was set at 7 ppm for parent mass and 20 ppm for fragment masses.

### PPI and Pathway Overrepresentation/Enrichment Analysis

Normalized protein signal was averaged across each condition and compared to the detection in corresponding IgG and knockdown controls. For HEK samples, proteins with 2-fold enrichment in CC2D1A IP samples or absent in their respective IgG controls were considered as candidate binding partners, then the hits shared across all three antibodies were chosen. For hippocampal samples, we selected targets that were significantly enriched by at least two-fold over IgG in the first experimental replicate, while the second replicate also required enrichment or absence in knockdown (1aKD) samples. Proteins that were shared between both hippocampal experiments were chosen. Candidate binding partners underwent functional over-representation/enrichment analysis using g:Profiler (https://biit.cs.ut.ee/gprofiler/gost) GO: Biological Process (GO:BP) and Cellular Compartment (GO:CC) with *Homo sapiens* or *Mus musculus* when experimentally appropriate for species and otherwise default parameters (37). PPI network analysis was conducted using all known interactions from the STRING database (https://string-db.org) and visualized with Cytoscape 3.10.3 (https://cytoscape.org) (38, 39). The full STRING network was built using 0.4 and 0.5 confidence cutoffs for the HEK293 and mouse IP networks, respectively. Proteins without any direct network connections (singletons) were then hidden. The STRING enrichments were conducted using a 0.5 overlap cutoff between GO:BP terms for both networks. For human interactors, additional analyses were performed using ShinyGO v0.82 (https://bioinformatics.sdstate.edu/go/) for KEGG pathway enrichment and eDGAR (http://ed-gar.biocomp.unibo.it/gene_disease_db/) for identification of genes associated with human disorders (40, 41). For SynGO (https://www.syngoportal.org) analysis of the mouse hippocampal interactors annotation was performed using gene counts for GO:CC terms including child terms (42).

### Subcellular Fractionation

Four freshly dissected hippocampi from two adult mice were lysed on ice using a glass dounce tissue grinder (Milipore Sigma:D8938) with 150 µL of lysis buffer prepared with 10 mM HEPES (2-(4-(2-hydroxyethyl) piperazin-1-yl) ethanesulfonic acid) pH 7.4 and 2 mM EDTA with protease and phosphatase inhibitors in sterile water. The lysate was centrifuged at 4°C for 5 minutes with 900 g. The homogenate fraction was collected from the supernatant for later analysis. The remaining sample was recentrifuged at 4°C for 15 minutes at 10,000 g to obtain the cytosolic light membrane fraction which was again collected from the supernatant. The pellet was resuspended in 100 µL of buffer containing 50 mM HEPES pH 7.4, 2 mM EDTA, 2 mM EGTA (ethylene glycol tetra acetic acid), 1% Triton-X-100 (VWR:97063-864) and protease and phosphatase inhibitors (both at 1:100 dilution) in sterile water and a portion was collected for the synaptosome fraction. The remaining synaptosome fraction was centrifuged at 4°C for 80 minutes at 10,000 g, resulting in the supernatant vesicular/presynaptic fraction. Finally, for the postsynaptic fraction, the pellet was resuspended in 30 µL 100 mM Tris pH 9 buffer containing 1% sodium deoxycholate, 0.2% sodium dodecyl sulfate, and 1:100 diluted protease and phosphatase inhibitors.

### Western Blot

For IP validation, 30 µL of protein lysate (i.e., full eluate) was separated by gel electrophoresis on 8% or 4-12% gradient bis-tris polyacrylamide gels (ThermoFisher:NW00082BOX and NW04120BOX, respectively) transferred in 1X Bio-Rad Tris/Glysine Buffer for western blots (Bio-Rad: 1610771) with 20% methanol onto Licor Odyssey Nitrocellulose Membrane (Licor:926-31092) for appropriate amounts of time given the gel concentration used. The membranes were incubated with the appropriate blocking agent (5% Milk in TBS-T, 3% BSA in TBS or 5% BSA in TBS with 0.1% Tween) for proteins of interest for 1 hour at room temperature with gentle rocking. Membranes were incubated with primary antibodies: mouse anti-CC2D1A 1:1000 (Abcam:ab68302), rabbit anti-CC2D1A 1:1000 (Abcam:ab191472), anti-CC2D1B 1:1000 (Proteintech:20774-1-AP), anti-TNIK 1:800 (Cell Signaling Tech: 32712S) for 24-48 hours at 4°C with gentle rocking.

For subcellular fractionation analysis, 10 mg of protein from each fraction was separated and blocked as above. Then, the membranes were probed for anti-CC2D1A 1:1000 (Abcam:ab68302), anti-CC2D1B 1:1000 (Proteintech:20774-1-AP), anti-PSD95 1:1000 (Neuromab:75028), and anti-Synaptophysin 1:10,000 (Abcam:32127) for 24-48 hours at 4°C under gentle rocking. Signal was normalized against the Revert™ 700 Total Protein Stain (Licor:926-11016). The secondary antibodies IRDye® 680 goat anti-mouse IgG (Licor:926-68020) and IRDye® 800CW goat anti-rabbit IgG (Licor:926-32211) were added at 1:10,000 and incubated for 1 hour at room temperature with gentle rocking. Blots were scanned using LI-COR Odyssey CLx and analyzed with Licor ImageStudioLite software (V5.2.5, 2015).

## RESULTS

### Comparative Analysis of CC2D1A Binding Partners Across Antibodies

Immunoprecipitation Mass Spectrometry (IP-MS) analysis has been a widely applied method to identify protein-protein interactions (PPI) for a given protein. However, IP efficiency remains highly protocol and antibody-dependent, which can lead to increased occurrences of type 1 errors (43). Since previous research on the CC2D1A interactome relied on an antibody showing substantial non-specific binding patterns (1), we compared enriched proteins from HEK293 lysates identified with IP-MS across three different commercial anti-CC2D1A antibodies: a rabbit polyclonal antibody used in the previous IP study targeting the N-terminal 50 amino acids of human CC2D1A (12), a mouse polyclonal raised against the full-length human protein (2, 44, 45), and a more recent rabbit monoclonal with a proprietary antigen (46). After filtering for false discovery peptides and comparing the data set to that of the appropriate IgG isotype control (**Fig. 1A-B**), we found that the rabbit monoclonal anti-CC2D1A antibody resulted in the highest enrichment of CC2D1A compared to the mouse and rabbit polyclonal antibodies (**Fig. 1C, Suppl. Table S1**). The enriched proteins from this data set showed notable overlap with those enriched by the two other antibodies (**Fig. 1D, Suppl. Table S1**). To define a high-confidence CC2D1A interactome, we focused on proteins that were enriched by at least two-fold in the CC2D1A IP compared to IgG control in all three antibodies or completely absent from the IgG control in one or more antibodies, identifying 208 proteins (**Suppl. Table S2**). Charged Multivesicular Body Protein 4B (CHMP4B) which had been previously validated at both the biochemical and ultrastructural level was present in all three IPs (16, 17). CHMP4B is a member of the ESCRT-III complex that assembles to mediate membrane remodeling during membrane scission, endolysosomal maturation, cell division, and virus budding (14, 47-49). CC2D1A has been involved in all these functions through its interaction with CHMP4B (14, 16, 17). It is important to note that this could be a conservative interaction network due to the limited number of identifications with the mouse polyclonal antibody. The rabbit polyclonal antibody had been previously used for IP-MS and 9 out 15 interactors identified in RAW cells in that study were also found in our enriched list from HEK293 cells using the same antibody (**Suppl. Table 1**), but were not present in the other IPs (12).

**Figure 1.**
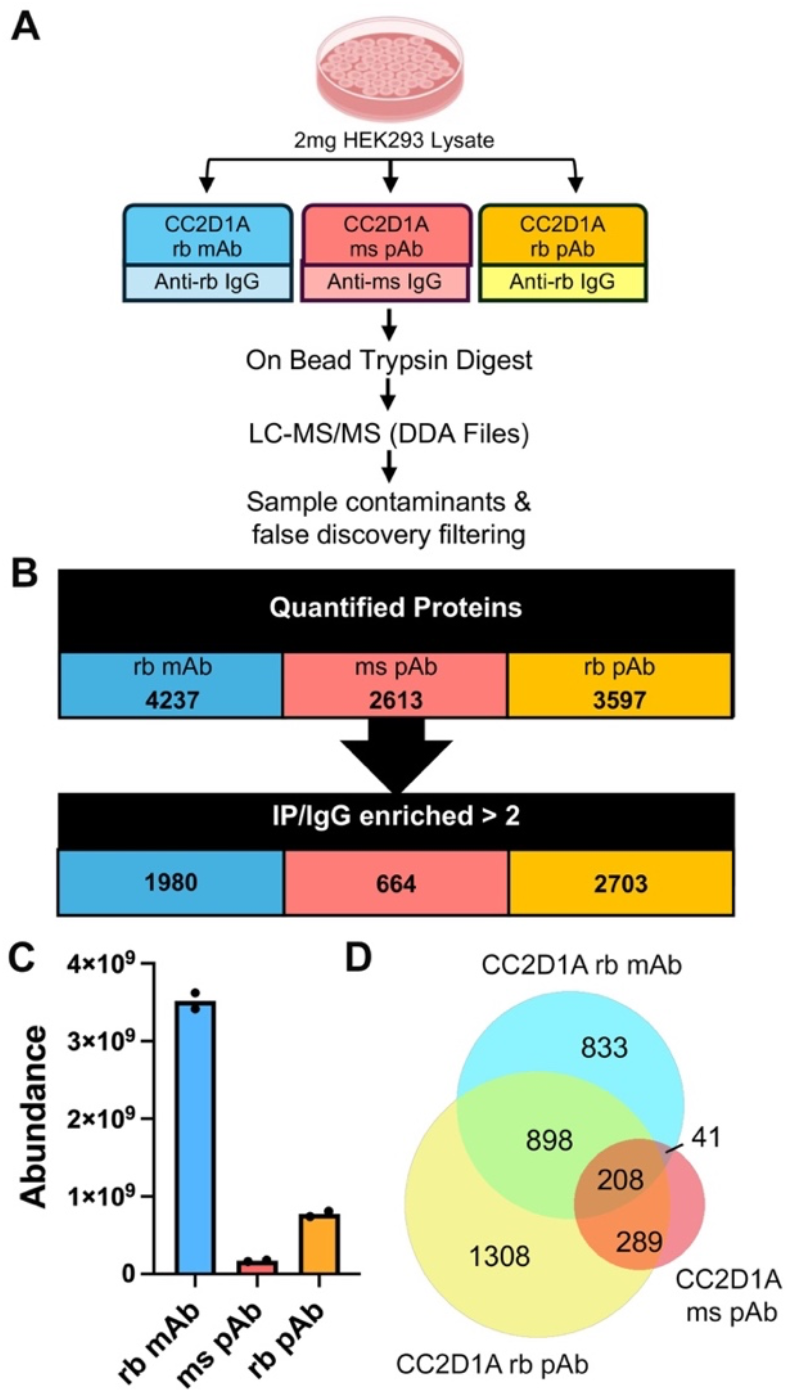
Experimental design for antibody validation and identification of the CC2D1A interactome in HEK293 cells. **A**, Workflow of endogenous CC2D1A complex pull-down across three commercial antibodies. Each anti-CC2D1A antibody (rabbit monoclonal: rb mAb, mouse polyclonal: ms pAb, rabbit polyclonal: rb pAb) and its appropriate isotype control were used on equal amounts of HEK293 lysate in duplicates and processed for LC-MS/MS. **B**, Filtering strategy. Total number of quantified proteins for both duplicates for each antibody were filtered for IP enrichment over IgG of greater than 2-fold. The list of enriched proteins for each antibody is available in **Suppl. Table 1**). **C**, Analysis of abundance (LFQ) values for CC2D1A indicated that the anti-CC2D1A rb mAb showed the greatest protein enrichment. **D**, Venn diagram showing the overlap in protein identification across the 3 antibodies. The 208 proteins that were shared across all 3 antibodies were chosen as the highest confidence interactors for follow-up analysis.

Gene ontology (GO) analysis via g:Profiler of the Cellular Component (GO:CC) associated with CC2D1A interactors showed strong enrichment for cytoplasmic proteins localized in organelles with 186 out of 208 linked with “cytoplasm” (GO:0005737) and 175 with “intracellular membrane-bounded organelle” (GO:0043231). The most significant representations within this group were in nucleoplasm (78 proteins), cytoplasmic vesicles (44 proteins), and mitochondria (33 proteins) with limited overlap between the sub-sets (**Fig. 2A**). Overlapping proteins mostly included kinases and proteasome components that could reside in multiple organelles. Interestingly, when we further probed the molecular function of nuclear proteins, they were primarily involved in RNA binding and protein binding. This suggests a regulatory role for CC2D1A in multiple subcellular locations.

**Figure 2.**
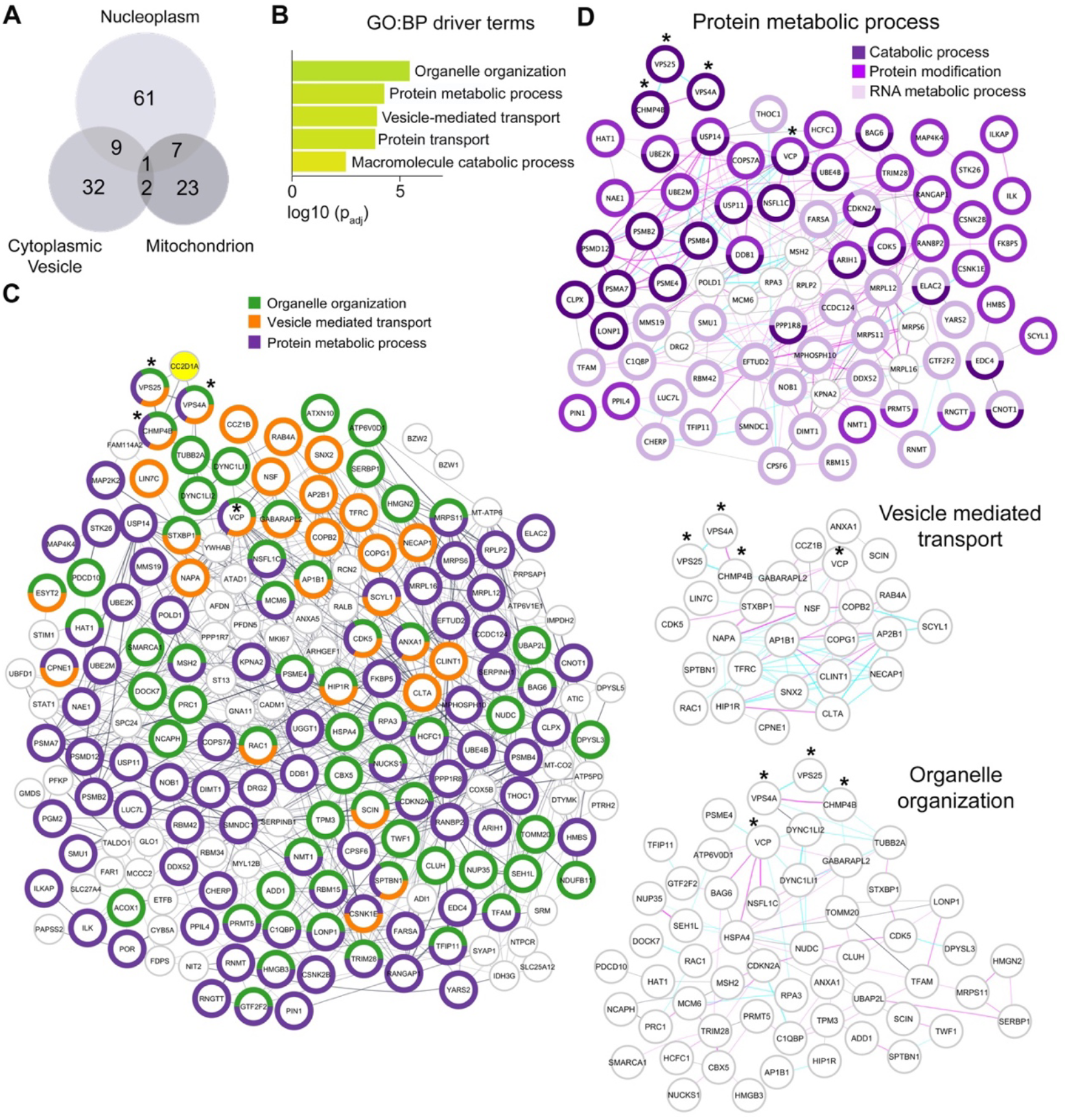
GO and PPI analysis of the CC2D1A binding complex in HEK293 cells. **A**, Venn diagram of protein cellular compartment overlap from GO:CC analysis performed in g:Profiler showing different subcellular localizations. **B**, Histogram of the driver biological process (GO:BP) terms identified pathways from GO:BP analysis. The full schematic with protein contributions to each term is in **Suppl. Fig.1. C**, STRING PPI network from confidence cutoff 0.4 and singletons hidden. Nodes represent individual proteins with network connections. Border color represents STRING functional enrichment GO:BP terms identified in g:Profiler. Edges represent various predicted and/or experimentally determined interactions between proteins, with edge thickness representing the strength of supporting data. **D**, Individual networks for organelle organization (GO:0006996), vesicle-mediated transport (GO:0016192), and protein metabolic process (GO:0019538) are shown on the right. Asterisks denote key CC2D1A interactors that are shared across all networks. Magenta edges denote experimentally determined interaction. Cyan edges denote database-suggested interaction. The protein metabolic process PPI network is further labeled for GO child terms highlighting specific functions in protein metabolism: catabolic process (GO:0009056), protein modification process (GO:0036211), and RNA metabolic process (GO:0016070).

Significantly enriched Biological Process GO terms (GO:BP) implicated CC2D1A in metabolic and protein trafficking functions such as “organelle organization” (GO:0006996), “protein metabolic process” (GO:0019538), and “vesicle-mediated transport” (GO:0016192) (**Fig. 2B, Suppl. Fig.2**). Functional enrichment analysis of the PPI network using STRING identified similar GO:BP terms, further supporting this regulatory role of CC2D1A (**Fig. 2C**). Analysis of the proteins involved in protein metabolism indicated a general effect on protein homeostasis, as proteins were split across RNA metabolism, protein modification including both ubiquitin modification and phosphorylation, and proteolysis with multiple core proteasome components represented including PSMA7, PSMB2, PSMB4, and PSMD12 (**Fig. 2C**). The closest interactors to CC2D1A in the STRING network are associated with all identified processes linking vesicle transport to protein homeostasis: ESCRT-III components CHMP4B and VPS4A, ESCRT-II component VPS25, and VCP, an ATPase involved in multiple aspects of organelle biogenesis and protein degradation (50). The loss of CC2D1A in murine and fly models has often shown deficits in proteostasis and endolysosomal biogenesis and these novel interactors suggest that CC2D1A may be deployed with CHMP4B to control different cellular functions (9, 17, 20, 51, 52).

Even if this dataset was generated in HEK293 cells, parallel KEGG pathway enrichment analysis using ShinyGO also identified “Pathways of neurodegeneration” as an enriched pathway due to multiple proteins involved in proteasome function and oxidative phosphorylation. Multiple among these novel interacting partners have been implicated in neurological and psychiatric diseases such as epileptic encephalopathy (STXBP1, DOCK7, NECAP1, SLC25A12, RANBP2), brain malformations (TUBB2A, CDK5, PDCD10), non-syndromic ID (HCFC1), neurodegenerative syndromes (ATXN10, SCYL1, VCP, MT-ATP6) (53-57). Additionally, the CC2D1A paralog CC2D1B was also identified as a candidate interactor in two of the three antibodies tested (**Suppl. Table 1**). With the cross-comparison of multiple primary antibodies for immunoprecipitation, we were able to identify high confidence-binding partners supporting a multifunctional role for CC2D1A in organelle stability and protein trafficking across various subcellular locations.

### The CC2D1A Interactome in Murine Brain Identifies a Synaptic Role

Null variants in *CC2D1A* in humans lead primarily to ID with comorbid ASD (1-5, 32). Thus, we investigated the CC2D1A interactome in the mouse hippocampus to identify interacting proteins that may be contributing to behavioral and intracellular signaling phenotypes. We used the rabbit monoclonal anti-CC2D1A antibody that was found to provide the greatest enrichment in the previous analysis (**Fig.1C**) and performed two replicate experiments on hippocampal lysates pooled from 2-3 mice per sample. In replicate 1, we performed four IPs (2 males and 2 females) with their respective IgG controls. In replicate 2, we additionally compared three hippocampal lysates from *Cc2d1a* hypomorph mice with 84% global reduction in CC2D1A protein expression (1aKD; *Cc2d1a*^*VH/VH*^ line;1 male and 2 females) with the WT sample to identify proteins that were significantly enriched against both the IgG and the hypomorph (**Fig.2A, Suppl. Fig. 1**). Candidate interactors were identified by filtering for proteins significantly enriched more than two-fold in the CC2D1A IP versus IgG control as well as significantly enriched versus the 1aKD line. Since many of the identified proteins do not have antibodies suitable for IP analysis, the second replicate would also provide independent validation of our initial results. Fourteen proteins were identified in both replicates and enriched against 1aKD in replicate 2 or only present in the CC2D1A IP. Two of them (RNF40 and SNRP) were removed because they were present in only 1 or 2 samples per replicate at low abundance resulting in a conservative group of 11 high-confidence interactors in the brain (**Fig. 3B, Suppl. Table S3**).

**Figure 3.**
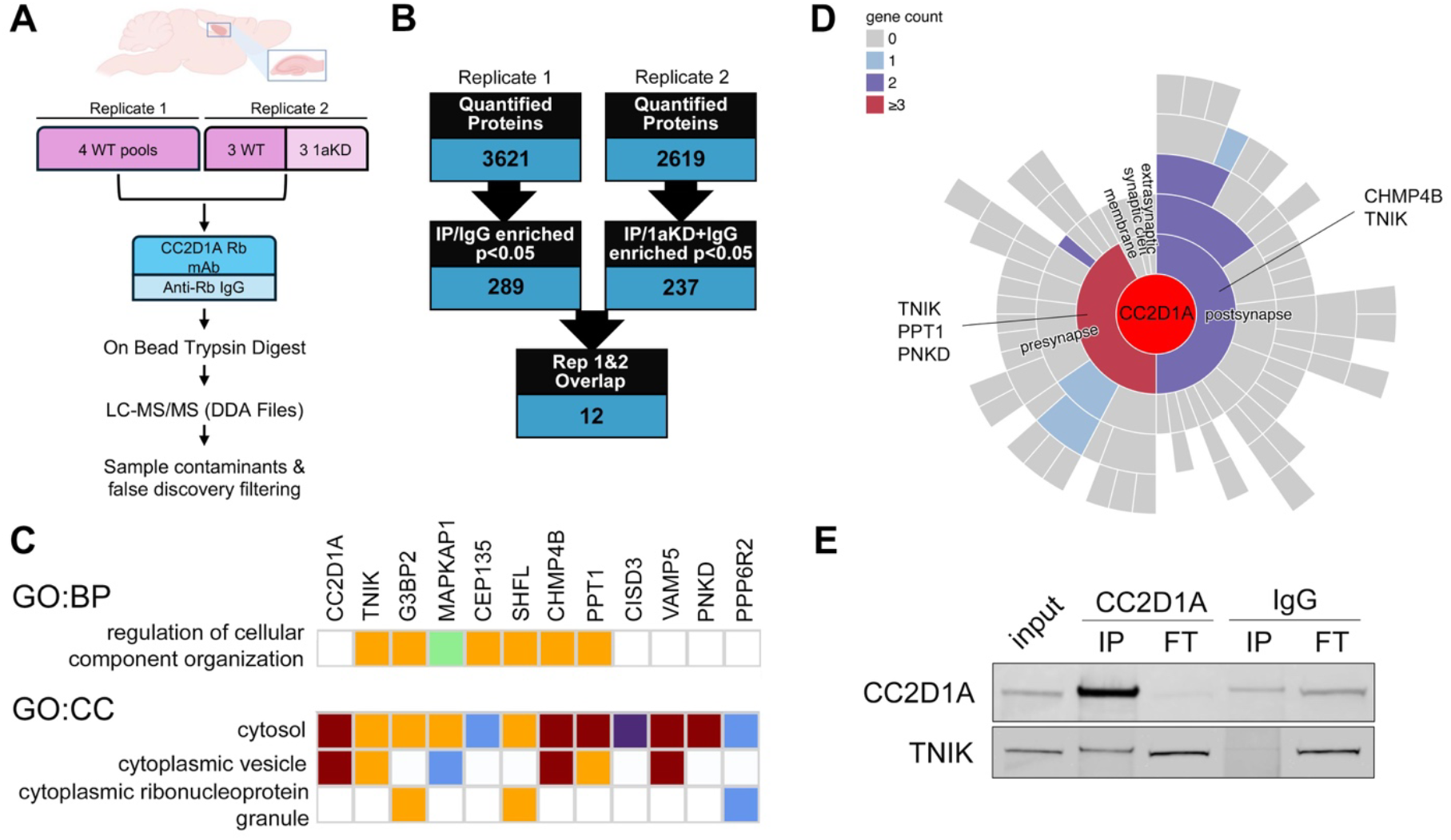
Identification of the CC2D1A interactome in mouse hippocampus. **A**, Workflow of endogenous CC2D1A pull down. WT and *Cc2d1a* knockdown (1aKD) mouse hippocampal lysate was purified with the rabbit anti-CC2D1A monoclonal antibody (Rb mAb) and anti-rabbit IgG control, trypsin digested and analyzed with LC-MS/MS. **B**, Filtering strategy. Quantified proteins in the CC2D1A IP were first filtered against the matching IgG control and then against the 1aKD samples to identify high confidence, bonafide interactors. The list of filtered proteins is available in **Suppl. Table 3**). **C**, g:Profiler annotations for enriched for GO:BP and GO:CC terms. Colors represent levels of evidence: Maroon, IDA – direct assay; Orange, ISO or ISS – sequence orthology or sequence similarity; Blue, IEA – electronic annotation; Purple, HDA – high-throughput direct assay; Green, NAS – non-traceable author. **D**, SynGO GO:CC analysis shows pre- and post-synaptic localization for CC2D1A interactors including CHMP4B. **E**, Immunoprecipitation (IP) analysis with CC2D1A antibody or IgG controls shows that a subset of TNIK interacts with CC2D1A in the mouse hippocampus (FT: flow through).

CHMP4B again emerged as an interactor and despite the small number of proteins, GO analysis still identified significantly enriched terms showing localization and function consistent with findings in the HEK293 interactome (**Fig.3C**). “Regulation of cellular component organization” (GO:0051128) was enriched as a Biological Process (p_adj_= 0.0394) and associated with 7 out of 12 proteins (TNIK, G3BP2, MAPKAP1, CEP135, SHFL, CHMP4B, and PPT1). Cellular Component terms, in addition to a cytoplasmic localization (GO:0005737 “cytoplasm”, p_adj_=0.00454), identified two independent groups with associations to cytoplasmic vesicles (GO:0031410, p_adj_=0.0148 for CHMP4B, TNIK, PPT1, MAPKAP1, and VAMP5, and to cytoplasmic ribonucleoprotein granules (GO:0036464, p_adj_=0.00969) for G3BP2, SHFL, and PPP6R2. These results are consistent with enriched functions in the broader HEK interactome in vesicle biogenesis and transport and RNA metabolism. The only overlapping proteins with the HEK293 identifications were CHMP4B and G3BP2, though G3BP2 was absent in the mouse polyclonal IP which had the lowest efficiency and lowest number of hits. Since *Cc2d1a* conditional removal from mouse forebrain results in male-specific cognitive and intracellular signaling deficits (29, 58, 59), we wondered whether there would be any difference in complex composition in males and females. We compared abundance of high confidence interactors between male and female samples and found no differences (**Suppl. Table 3**).

As CC2D1A and CHMP4B are both localized at synapses, we reanalyzed the protein list using SynGO, a curated annotation tool that focused specifically on synaptic proteins (42). In addition to CC2D1A, 4 additional proteins were identified as synaptic proteins with CHMP4B and TNIK at postsynaptic specialization and PPT1, PDNK, and again TNIK at the presynapse (**Fig.3D**). While loss of *Cc2d1a* in mouse models impairs the maintenance of long-term potentiation in the hippocampus (29, 60), previous studies have found no changes in pre-synaptic release in culture or in slices (28, 60). TNIK (Traf2 and NcK interacting kinase) is highly enriched at the postsynaptic density and is a key factor in maintaining synaptic composition and stability functioning as a signaling hub (61, 62). Biallelic loss of function variants in *TNIK* also lead to intellectual disability (63). Due to the role of CC2D1A as a regulator of diverse signaling pathways, we sought to validate the interaction with TNIK via immunoblot. We found that a subset of TNIK interacts with CC2D1A, thus identifying a novel signaling interactor at the post synapse (**Fig.3E**).

### Differential Localization of CC2D1A and CC2D1B at Synapses

In investigating the synaptic role of CC2D1A we asked about the relationship with CC2D1B. Even though CC2D1B was not identified in the brain interactome, its identification in HEK293 cells spurred us to further investigate its role in neurons due to the strong evidence on functional redundancy between the two paralogs. CC2D1A and CC2D1B have similar domain structure with differences in the localization of the coiled-coil domains (**Fig. 4A**), but only CC2D1A loss of function has been reported to cause neurodevelopmental disorders in humans (1, 2). While loss of the only CC2D1 fly ortholog, lethal (2) giant discs (Lgd), results in larval lethality, expression of either human CC2D1A or CC2D1B can rescue this phenotype, suggesting functional redundancy (52). Similarly, *Cc2d1a* and *Cc2d1b* double knock-out embryos die before E11.5 and a single *Cc2d1a* or *Cc2d1b* allele can support embryonic development in the absence of the other gene (58). However, behavioral analyses revealed a more complex interplay in the brain. Conditional removal of *Cc2d1a* in the forebrain leads to cognitive and social deficits and hyperactivity, while *Cc2d1b* KOs only show cognitive deficits (29, 58). However, double heterozygotes are more similar to *Cc2d1a* conditional KO mice suggesting that loss of *Cc2d1a* may have more severe impacts on brain function (58).

**Figure 4.**
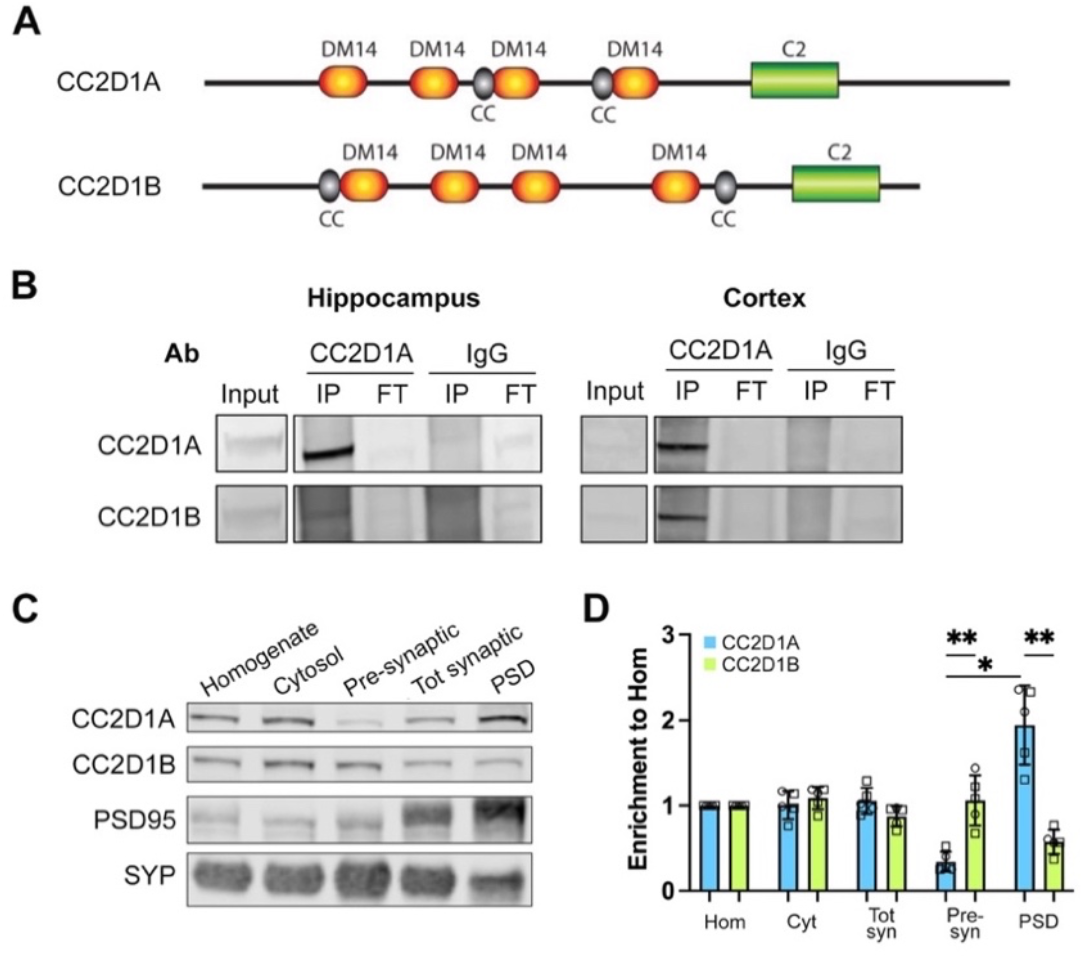
CC2D1A can bind to CC2D1B but is preferentially localized to the PSD. **A**, Protein schematic showing the domain structure of mouse CC2D1A and its paralog CC2D1B. **B**, Representative images of CC2D1A adult hippocampus and cortex IP probed for CC2D1A and CC2D1B. **C**, Representative image of subcellular fractionations for CC2D1A and CC2D1B. PSD95 and synaptophysin (SYP) are used to indicate post and presynaptic enrichment, respectively. **D**, Quantification of ratios for CC2D1A and CC2D1B in subcellular fractions compared to the homogenate (Hom). Data are presented as mean ± SEM showing significant multiple comparisons for the presynaptic (Pre-syn) and PSD fractions. * p<0.05, ** p<0.01, adjusted p-value. Cyt: cytosol, Tot syn: total synaptic. Three male (□) and two female (◯) WT samples were used.

Our findings in HEK cells indicated that the CC2D1 proteins can function in a complex. When we tested whether anti-CC2D1A antibodies could pull down CC2D1B in the brain, we found that it was present in both hippocampal and cortical lysates (**Fig. 4B**). A previous study showed a possible postsynaptic enrichment of CC2D1A (60). Thus, we performed synaptosomal fractionation to obtain fractions enriched for pre- and postsynaptic proteins in multiple samples to quantify these differences and tested for CC2D1B protein distribution in these fractions. We found significant differences in CC2D1A and CC2D1B localization (**Fig. 4C**). 2-way ANOVA showed a significant effect of protein (p=0.029 *) and fraction (p=0.0016 **) and of protein/fraction interaction (p<0.0001 ***). Multiple comparisons identified significant enrichment of CC2D1A in the PSD vs. pre-synaptic fractions (p=0.0204 *) and significant differences between CC2D1A and CC2D1B at the presynaptic (p=0.0034 **) and PSD fractions (p=0.0018 **) (**Fig. 4D**). This result indicates that CC2D1A may have a CC2D1B-independent function at the PSD underlying the more severe behavioral deficits in mice and ID in humans.

## DISCUSSION

This study identified high-confidence interactors for CC2D1A in both adult mouse hippocampus as well as a human cell line (HEK293). To date, this represents the most comprehensively curated set of proteins that either directly or indirectly bind to CC2D1A, analyzed across three different commercially available antibodies which target CC2D1A with varying levels of specificity. These interactions are endogenous, with the brain interactors verified against a knockdown control (1aKD) and the selection criteria were conservative to limit the identification of false positives.

CC2D1A had been involved in a variety of cellular functions ranging from endosomal trafficking, intracellular signaling in multiple pathways in cancer, immunity, and synaptic plasticity (12-14, 18, 20). Determining a unifying function had been confusing as multiple studies relied on two-hybrid assays or overexpression. The endogenous CC2D1A interactors we identified in HEK cells with three independent antibodies support a role as a multifunctional protein scaffold that can act at different subcellular locations. We found that CC2D1A interactors may independently localize in the nucleus, mitochondrion, and cytoplasmic vesicles but share functions in protein metabolism and organelle organization. These interactions converge on a set of four common interactors (CHMP4B, VPS4A, VPS25, and VCP) that are shared across the different functional modules. CHMP4B, VPS4, and VPS25 are all components of the ESCRT complex which contributes to a multitude of cellular events such as endolysosomal transport and multivesicular body formation (48), cytokinetic abscission (49), retroviral budding (47), microvesicle shedding, as well as cilia formation (64, 65). The AAA+-type ATPase VPS4 disassembles the membrane-bound ESCRT-III filaments (50), while VPS25 is part of the ESCRT-II complex that regulates both ESCRT-III assembly and capture of ubiquitinated proteins for trafficking (66). These identifications support the critical role of CC2D1A as an integral regulator of ESCRT function wherever the complex is acting.

In parallel, VCP, also known as p97, is AAA+-type ATPase that can link endosomal trafficking to protein homeostasis. VCP has a critical role in protein-quality control as it can bind ubiquitinated proteins and prime them for degradation and has a general role in organelle homeostasis (67). The VCP interactome is tightly linked to our CC2D1A interactome involved in protein modification including both the proteasome complex and multiple ubiquitin ligases (68). Multiple other CC2D1A interactors support a role in protein trafficking and proteostasis and a link to the signaling endosome. An elegant series of studies showed how CC2D1A is involved in endosomal trafficking and signaling of endosomal Toll-Like Receptors (TLRs) and RIG-I during immune responses (19, 20, 59). Loss of function of the CC2D1 *Drosophila* ortholog Lgd leads to dysregulated endolysosomal transport where late endosomes fail to fuse with lysosomes resulting in signaling proteins such as Notch and BMP/Dpp to accumulate without being degraded and sustain signaling (52, 69, 70). Overall, the HEK interactome indicates CC2D1A is involved in protein trafficking and degradation via the endolysosomal pathway, but it can be deployed as needed where the ESCRT complex functions.

This association with the diverse ESCRT functions and signaling is reflected in the hippocampal interactome which reprises the organelle organization role linking CC2D1A to synaptic function. With CHMP4B as the shared thread to ESCRT function, neuronal interactors are enriched for localization in cytoplasmic vesicles (TNIK, MAPKAP1, PPT1, VAMP5), but also reveal a possible role in cytoplasmic RNA granules (G3BP2, SHFL, PPP6R2). Both G3BP2 and SHFL are RNA binding proteins and G3BP2 was also identified in two different HEK IPs (rabbit polyclonal and monoclonal Abs) (**Suppl. Table 1**). Cytoplasmic RNA granules store and transport RNAs to control localization of translation and have a critical role in neuronal plasticity (71). PPT1 and PNKD are presynaptic proteins involved in neurotransmitter release and mutation lead to neurological conditions, Infantile Neuronal Ceroid Lipofuscinosis and familial Paroxysomal Nonkinesigenic Dyskinesia, respectively (72, 73). However, no synaptic vesicle release deficits have been identified in *Cc2d1a* KO mice in multiple studies (28, 60). Previous work showed substantial functional redundancy between the two CC2D1 proteins as either one could rescue mortality in Drosophila and mouse models and regulate ESCRT polymerization (52, 58, 74). Yet, there was a functional divergence, as *Cc2d1b* KO mice have milder cognitive deficits and *CC2D1B* has yet to be associated with human disease. Our data indicate that CC2D1B is enriched at the presynapse compared to CC2D1A and it is possible that CC2D1B could compensate for loss of CC2D1A. We propose that CC2D1A and CC2D1B can act in parallel or independently, and it is likely that loss of CC2D1A has more severe effects on cognitive function due to a preferential post-synaptic function. The relative contribution of the CC2D1 proteins to synaptic transmission deserves further investigation.

Deficits in *Cc2d1a* KO mice have pointed to a primarily postsynaptic role regulating synapse number and long-term potentiation (LTP) (2, 28, 60). Among our interactors, CHMP4B is enriched in PSDs, as are other ESCRT-III proteins, where they are thought to regulate membrane trafficking (75). ESCRT function at the PSD is still not well understood and CHMP4B has also been involved in dendritic branching in *Drosophila* where it is termed *shrub* because its loss increases branching (75-77). Previous work also attributed synaptic plasticity deficits caused by loss of CC2D1A to disruption in different signaling pathways. Increased activity of RAC1, a Rho GTPase involved in actin polymerization, was involved in spine density and late LTP deficits that could be rescued with Rho inhibition (60). Other studies identified an interaction with PDE4D, that also causes hyperactivation upon loss of CC2D1A and a reduction in cyclic AMP (cAMP) signaling (12, 23). Similarly, inhibition of PDE4D activity can rescue behavioral and signaling deficits in *Cc2d1a*-deficient mice (23). Both RAC1 and PDE4D were identified in HEK IPs and hippocampal replicate 1, but they were not enriched over IgG control in the brain samples. If these interactions are present endogenously in the brain, they may be transient and affecting only a subset of the protein. This was evident in our validation IP for TNIK where most of the protein was still in the flowthrough and only a portion was pulled down with CC2D1A. TNIK is a MAP kinase kinase kinase kinase (MAPK4) mutated in intellectual disability (63). TNIK is also enriched at the PSD and has been shown to regulate the stability of PSD components and synaptic maturation (61, 62, 78). Modulating TNIK activity could represent an additional target to rescue phenotypes caused by loss of CC2D1A focusing specifically on the synapse since both RAC1 and PDE4D are expressed in multiple tissues.

Overall, results from our interactome analysis in two independent cell/tissue types support a pleiotropic role for CC2D1A in regulating ESCRT function in different subcellular organelles reconciling the different roles identified in the previous literature. We also revealed how interactions with synaptic proteins and CC2D1B could underlie the synaptic plasticity and neurodevelopmental deficits in *CC2D1A*-related disorders.

## Acknowledgements

We would like to thank the Rutgers Proteomic Facility and in particular Caifeng Zhao with assistance with these experiments, and Peter Nemes (University of Maryland), Sila Ultanir (The Francis Crick Institute), Anurag Raj (University of Pittsburgh), Dhirendra Kumar (Rancho BioSciences/ Johnson & Johnson), Amit Yadav (Translational Health Science and Technology Institute, India) for comments on the manuscript and data analysis.

## Author contributions

A.T.H., A.B., L.T. and M.C.M. designed the research; A.T.H., A.B., L.T., S.T., E.S., A.M., and H.Z. conducted experiments; A.T.H., A.B., H. L., L.T., and M.C.M. analyzed the data; A.T.H., A.B., and M.C.M. wrote the manuscript; all authors were involved in manuscript review and editing.

This article contains supplemental data.

