## Supplemental Information for "Interactome Analysis of the CC2D1A Scaffold Reveals Novel Neuronal Interactions and a Postsynaptic Role"

**Supplementary Figures 1 and 2**

**Legends for Supplementary Tables 1-3**

### Supplementary Figures

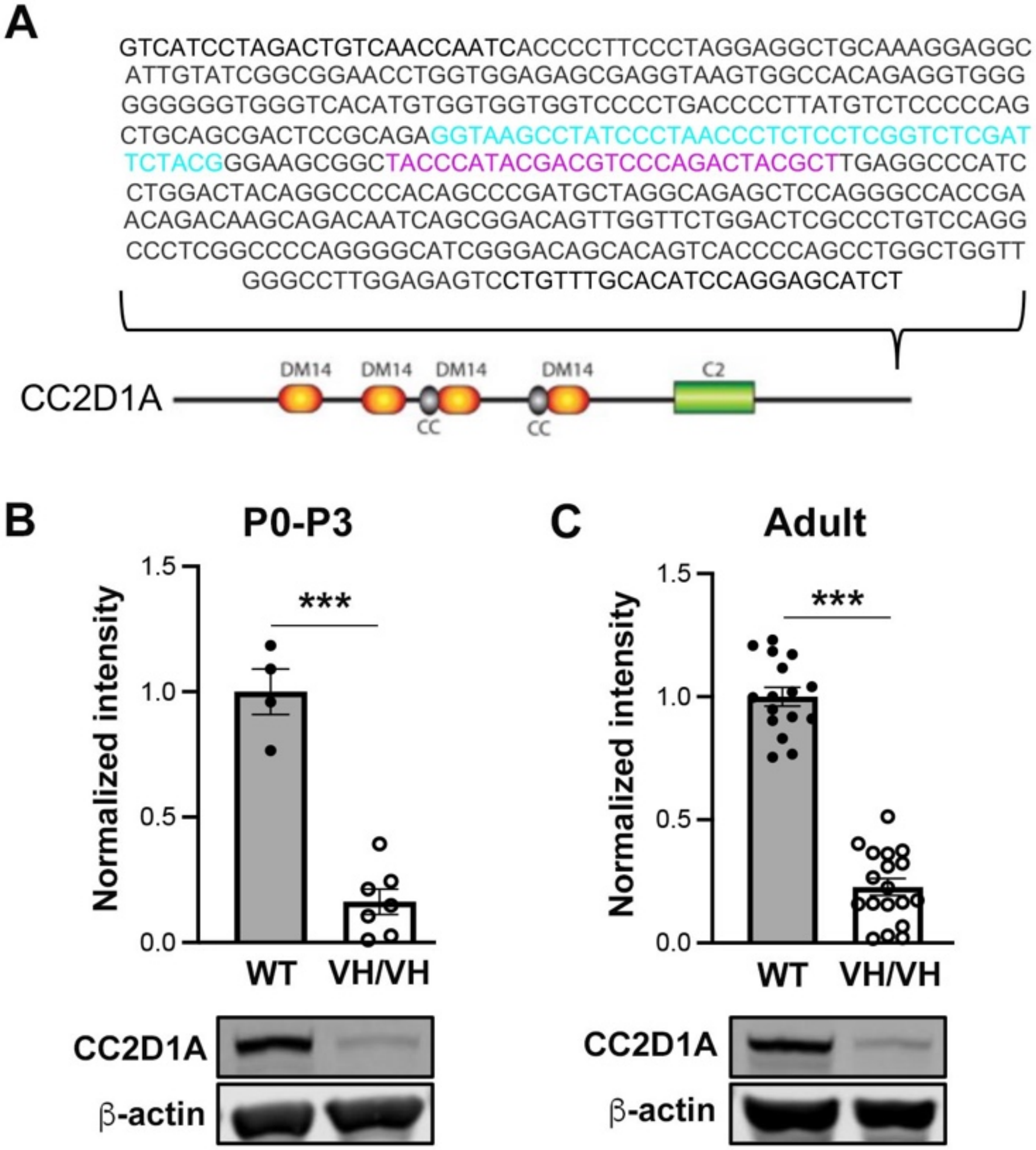


**Supplementary Figure 1.** ***Cc2d1a*$^{VH/VH}$ line**. ***A,*** The V5-HA line was created by inserting a 42bp V5 (cyan) and a 27bp HA (magenta) tags in the 3' end of the mouse *Cc2d1a* gene. ***B,*** Quantification of CC2D1A protein expression via Western blot analysis in the wild-type (WT) and *Cc2d1a*$^{VH/VH}$ P0-P3 hippocampi. **C,** Quantification in adult hippocampi probed for CC2D1A. Data are presented as CC2D1A intensity (mean ± SEM) normalized to endogenous β-actin control and then compared to the average of WT samples from each Western blot (*** p-value <0.001, t-test)

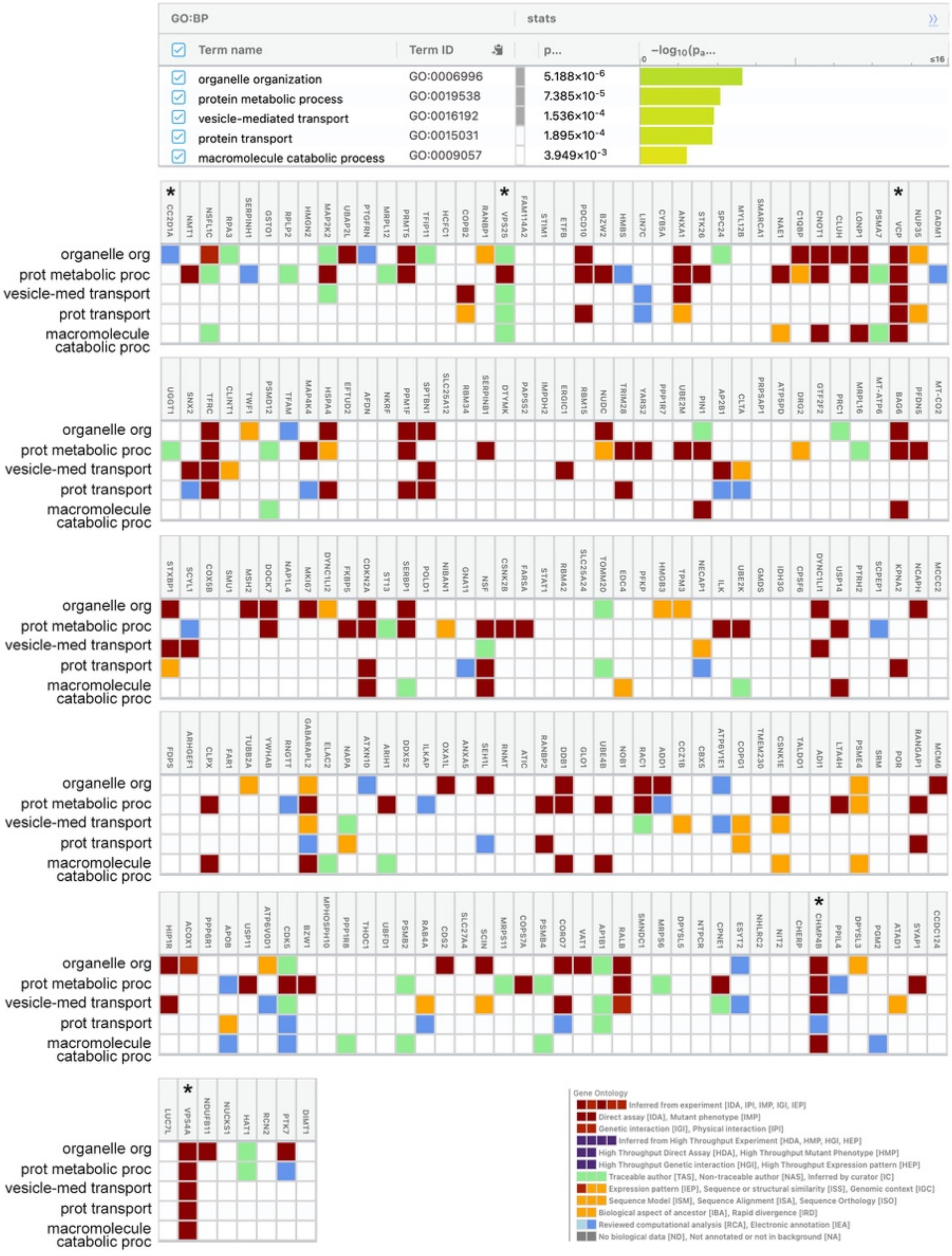


**Supplementary Figure 2. GO:BP term contribution for individual proteins in the CC2D1A interactome in HEK293 cells.** Asterisks denote that the proteins are also labeled in Figure 2C.

**Supplementary Table Legends**

**Supplementary Table 1:** Lists of quantified and enriched proteins for each of 3 different anti-CC2D1A antibodies (rabbit monoclonal, mouse polyclonal, and rabbit polyclonal antibodies). Index in first sheet.

**Supplementary Table 2:** GO analysis results for 208 candidate CC2D1A interactors in HEK293 cells

**Supplementary Table 3:** Lists of quantified and enriched proteins for the 2 replicate experiments using anti-CC2D1A IP in the mouse hippocampus. Summary of high confidence hits and comparison of their abundance between male and female mice. Index in first sheet.
